# NanoVar: Accurate Characterization of Patients’ Genomic Structural Variants Using Low-Depth Nanopore Sequencing

**DOI:** 10.1101/662940

**Authors:** Cheng Yong Tham, Roberto Tirado-Magallanes, Yufen Goh, Melissa J. Fullwood, Bryan T.H. Koh, Wilson Wang, Chin Hin Ng, Wee Joo Chng, Alexandre Thiery, Daniel G. Tenen, Touati Benoukraf

**Affiliations:** Cancer Science Institute of Singapore, National University of Singapore, Singapore 117599, Singapore; School of Biological Sciences, Nanyang Technological University, Singapore 637551, Singapore; Department of Orthopedic Surgery, National University Health Systems, Singapore 119228, Singapore; Department of Orthopaedic Surgery, Yong Loo Lin School of Medicine, National University of Singapore, Singapore 119228, Singapore; Department of Hematology-Oncology, National University Cancer Institute of Singapore, National University Health System, Singapore 119228, Singapore; Department of Medicine, Yong Loo Lin School of Medicine, National University of Singapore, Singapore 119228, Singapore; Department of Statistics and Applied Probability, National University of Singapore, Singapore 117546, Singapore; Harvard Stem Cell Institute, Harvard Medical School, Boston, MA 02115, USA; Discipline of Genetics, Faculty of Medicine, Memorial University of Newfoundland, St. John’s, NL, A1B 3V6, Canada

## Abstract

Despite the increasing relevance of structural variants (SV) in the development of many human diseases, progress in novel pathological SV discovery remains impeded, partly due to the challenges of accurate and routine SV characterization in patients. The recent advent of third-generation sequencing (3GS) technologies brings promise for better characterization of genomic aberrations by virtue of having longer reads. However, the applications of 3GS are restricted by their high sequencing error rates and low sequencing throughput. To overcome these limitations, we present NanoVar, an accurate, rapid and low-depth (4X) 3GS SV caller utilizing long-reads generated by Oxford Nanopore Technologies. NanoVar employs split-reads and hard-clipped reads for SV detection and utilizes a neural network classifier for true SV enrichment. In simulated data, NanoVar demonstrated the highest SV detection accuracy (F1 score = 0.91) amongst other long-read SV callers using 12 gigabases (4X) of sequencing data. In patient samples, besides the detection of genomic aberrations, NanoVar also uncovered many normal alternative sequences or alleles which were present in healthy individuals. The low sequencing depth requirements of NanoVar enable the use of Nanopore sequencing for accurate SV characterization at a lower sequencing cost, an approach compatible with clinical studies and large-scale SV-association research.

## Introduction

Structural variations are implicated in the development of many human diseases^1,2^ and account for most of the genetic variation in the human population^3,4^. Structural variants (SVs), defined as genomic alterations greater than 50 bp^5^, can functionally affect cellular physiology by forming genetic lesions which may lead to gene dysregulation or novel gene-fusions, driving the development of diseases such as cancer^6,7^, Mendelian disorders^8,9^, and complex diseases^10^. SVs can exist as different classes including deletion, duplication, insertion, inversion and translocation. Over the years, disease-associated SVs were indicated as biomarkers for diagnosis^9,11^, prognosis^12^, and therapy guidance for patients^13^, which could be screened through sequencing-based and non-sequencing-based methods in clinics. As the clinical impacts of SVs continue to unveil, there is a clear need for accurate, rapid and inexpensive workflows for routine SV profiling in patients to expedite biomarker discovery and broaden clinical investigations^7^.

There are currently two main standards of sequencing-based methods for comprehensive SV detection: long-read or third-generation sequencing (3GS) and short-read or second-generation sequencing (2GS). Although 3GS technologies were made accessible to a large audience, it has not yet supplanted 2GS technologies due to its higher sequencing error rate and lower throughput^14^. While 3GS is currently mainly restricted to the study of small genomes^15^ or targeted sequencing^16^, recent studies have reported mammalian whole-genome sequencing (WGS)^17,18^ but at a higher sequencing cost per megabase as compared to the older technologies. In the domain of SV discovery, many groups have reported that 3GS approaches provided higher SV detection sensitivity and resolution than 2GS, despite their higher sequencing error rate^9,11,19^. This is mostly due to the inadequacy of short reads (50-200 bp) to elucidate large genomic variations also involving novel sequence insertions or repetitive elements, which may give rise to high false discovery rates^5,20^. On the other hand, longer read lengths (>1 kb) reduce mapping ambiguity, resolve repetitive sequences^21^ and complex SVs^8^, and discover a much larger extent of SVs than short-reads^17,18^. Despite better SV detection capabilities, the low throughput and high sequencing cost per megabase of 3GS obstruct its feasibility to be used in routine SV interrogation in patients.

To overcome these issues, we developed a new SV caller tool, NanoVar, which utilizes low-depth Oxford Nanopore Technologies (ONT) WGS data for accurate SV characterization in patients. NanoVar adopts a neural-network-based algorithm for high-confident SV detection and SV zygosity estimation for all SV classes. It is optimized to work with shallow long-read WGS data at a minimum sequencing depth of 4X or 12 gigabases (Gb) in total bases, which can be achieved with one to five ONT MinION sequencing runs, depending on the flowcell chemistry, library preparation kit, and sample quality. In this manuscript, we evaluated NanoVar’s SV detection precision and recall amongst other tools using simulation datasets. When applied to patient data, we demonstrated the feasibility and speed of implementing the NanoVar workflow for SV discovery in low depth 3GS clinical samples.

## Results

### The NanoVar workflow

The NanoVar workflow is a series of processes that utilizes 3GS long reads to discover and characterize SVs in DNA samples. The sequencing of the genome of interest is carried out by ONT MinION to produce long-reads reported in FASTQ/FASTA formats (Fig. 1a). The sequencing output of several sequencing runs can be combined to achieve enough depth of coverage for SV discovery. Based on our initial tests performed in simulated dataset (*cf.* hereafter), we recommend having at least 12 Gb of sequencing data (covering approximately 4X of the human genome), which can be achieved through one to five MinION runs depending on the flowcell chemistry (R9.4, R9.5), library preparation kit (1D, 1D^2^, 2D), and DNA sample quality (purity, quantity and fragment length). The R9.4 flowcell chemistry with a 1D sequencing library using high purity DNA with a mean fragment length of 8 kb provides the highest throughput per MinION flowcell (above 12 Gb). The combined FASTQ/FASTA file is used as input into the NanoVar tool for SV processing. NanoVar begins by mapping the long reads against a reference genome using HS-BLASTN^22^ to obtain the alignment profile of each read (Fig. 1b). Reads with incomplete alignments (containing divergent sequence/gap) are selected and evaluated through an SV characterization algorithm (Supplementary Fig. 1) to characterize for the possible SV classes. NanoVar can distinguish six classes of SV: deletion (DEL), inversions (INV), tandem duplication (DUP), insertion (INS, novel sequence insertion/insertion of sequences absent from reference genome), genomic insertion (insertion of sequences found elsewhere in the reference genome) and translocation. Due to the close resemblance in altered sequences between a genomic insertion and a translocation, they are collectively labeled as “breakends” (BND). After all reads are classified, NanoVar calculates the read-depth coverages for all the SV breakend sites, separating the number of breakend-supporting reads and breakend-opposing reads. Lastly, the read-depth coverage of each SV, together with other SV characteristics, are used as features for a simulation-trained neural network classifier to determine a confidence score for each SV. This confidence score is used to rank the SVs by confidence and reduce false positives in the final output. The filtered list of SVs is recorded in a standard Variant Calling Format (VCF) file and an HTML report. The HTML report provides an overview of the SV analysis and an SV output table containing the information of each SV which can be filtered and downloaded in MS Excel or CSV formats. The figures presented in the report also include an SV class distribution chart and read length distribution of the sequencing reads which serves to QC for the input (Fig. 1c). NanoVar also assigns a breakend read ratio value to each SV to estimate their SV zygosity, where a ratio of 1.0 refers to a homozygous estimation and 0.5 refers to a heterozygous estimation.

**Figure 1:**
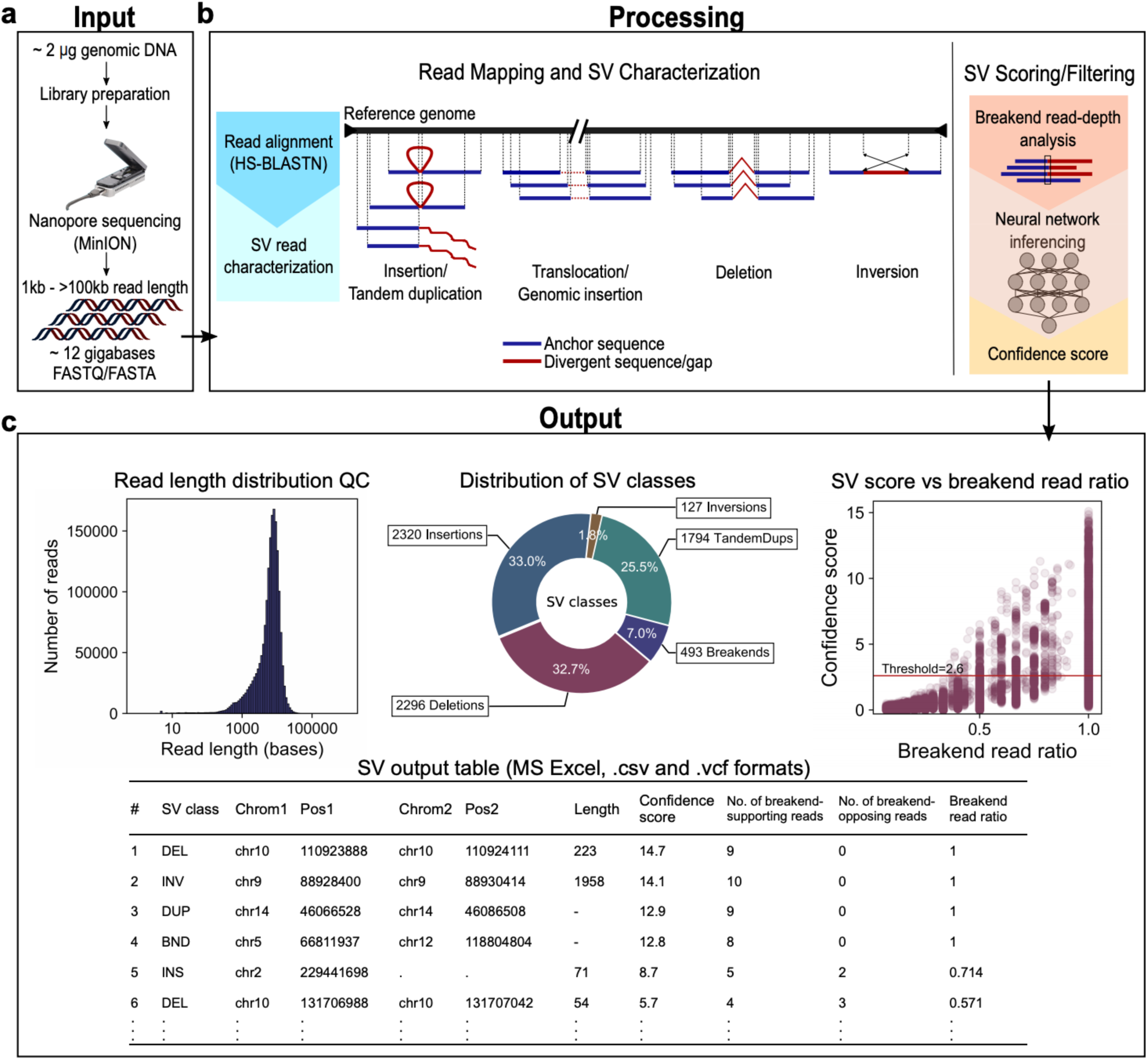
The NanoVar Workflow. (a) About 2 µg of human genomic DNA is set aside for library preparation and Nanopore sequencing to generate 3GS long sequencing reads. Long reads from all sequencing runs are combined into a single FASTQ/FASTA file (at least 12 gigabases) which is used as input into NanoVar. (b) NanoVar SV characterization process. (Left) Long reads are aligned to a reference genome using HS-BLASTN to identify anchor sequences (blue) and divergent sequences or gaps (red) within each read. Next, the alignment information is used to detect and characterize the different SV classes. (Right) For each characterized SV, read-depth coverage is calculated at their breakend(s) site for the number of breakend-supporting and breakend-opposing reads. The breakend read-depth, together with other alignment information, are employed as features in a neural network model to infer a confidence score for each SV. (c) NanoVar outputs all characterized SVs in a VCF file and produces an HTML report for QC and results visualization. The following figures can be found in the HTML report. (Top-left) Histogram showing the length distribution QC of the input sequencing reads. (Top-middle) Donut chart showing the distribution of SV classes characterized in the dataset (after confidence score filtering). Breakends represent translocation or genomic insertion SV. (Top-right) Scatter plot displaying the confidence score and breakend read ratio (fraction of breakend-supporting reads at a breakend) of each SV, also showing the confidence score threshold parameter used for filtering (red line). (Bottom) Table showing the details of all characterized SVs, which can be sorted, filtered and extracted in CSV or MS Excel formats.

### Benchmarking NanoVar using simulations

To evaluate NanoVar’s performance among other existing SV callers, we utilized three simulated datasets (doi:10.5281/zenodo.2599376). Each dataset contains about 1,150 randomly located SVs of different classes with sizes ranging from 25 bp to 100,000 bp (Supplementary Fig. 2a). Sequencing reads of both long-read and short-read sequencing were simulated to a sequencing depth of 4X and 50X respectively and were used as input for the workflows of each tool. Sniffles^19^, Picky^23^ and NanoSV^24^ are 3GS long-read SV callers while novoBreak^25^ and Delly^26^ are 2GS short-read SV callers. Taking the ground truth SVs in each simulation dataset as a reference, the precision and recall exhibited by each tool could be calculated judging from the genomic location of each SV, ignoring SV class annotation accuracy (Supplementary Table 1). Among long-read SV callers, NanoVar outperformed the rest in precision and recall, achieving the highest average F_1_ score of 0.91 for all three simulations (Fig. 2a). At the confidence score threshold of 1.4, NanoVar detected 85% of the ground truth with a high precision of 0.97, while the other long-read SV callers had slightly lower sensitivities (Sniffles: 0.81, NanoSV: 0.76, Picky: 0.81) and substantially lower precisions (Sniffles: 0.70, NanoSV: 0.21, Picky: 0.002), due to the reporting of more false positive SVs. Short-read SV callers generally exhibited higher precisions (Delly: 0.96, novoBreak: 0.99) and moderately lower recall (Delly: 0.78, novoBreak: 0.66) than long-read SV callers. To investigate the sensitivity of each tool in detecting SVs of different sizes, we examined the recall of each SV size and discovered that SVs with sizes ranging from 25 bp to 50 bp were inadequately detected by NanoVar and all other tools except Picky (Fig. 2b). After omitting these SVs from the datasets, we saw a significant improvement in the F_1_ scores of all tools except Picky, with an increased in NanoVar’s recall from 0.85 to 0.96 (Supplementary Fig. 3a). To investigate NanoVar’s capability to handle datasets of different sequencing depth coverages, we tested NanoVar on simulated datasets with 1X to 40X depth of coverages generated from simulation 1 (Supplementary Fig. 3b). At low coverages (1X and 2X), NanoVar performed inadequately with F_1_ scores between 0.70 and 0.90. With coverage of 4X, NanoVar was able to achieve an F_1_ score close to its maximum, which is similar for coverages higher than 4X (8X, 16X, 32X, 40X). These results imply that 4X is the minimal coverage required for optimal NanoVar SV characterization and that increasing the coverage above 4X will not affect its performance. To evaluate the SV class characterization accuracy of each tool, we calculated the F_1_ scores for each SV class separately considering the SV class annotation accuracy of each tool in simulation 1 with 25 bp and 50 bp SVs omitted (Fig. 2c, Supplementary Table 2). Short-read SV callers achieved high F_1_ scores (between 0.72 – 0.98) for all SV classes except INSs, of which none was detected for both tools. Conversely, long-read SV caller Sniffles was able to characterize all SV classes but with lower F_1_ scores (between 0.51 – 0.86). NanoSV was able to characterize INSs but was inadequate in characterizing other SV classes, mainly due to them having incorrect SV class annotations. Picky performed relatively well in characterizing DUPs, INVs and BNDs, but had acutely low F_1_ scores for DELs and INSs due to the high number of false positives. Lastly, NanoVar was able to characterize all SV classes with higher F_1_ scores than the other 3GS SV callers (DUP: 0.96, DEL: 0.98, INS: 0.62, BND: 0.90, INV: 0.90). In conclusion, NanoVar exhibits higher SV characterization recall and precision amongst other long-read SV callers at a sequencing coverage of 4X.

**Figure 2:**
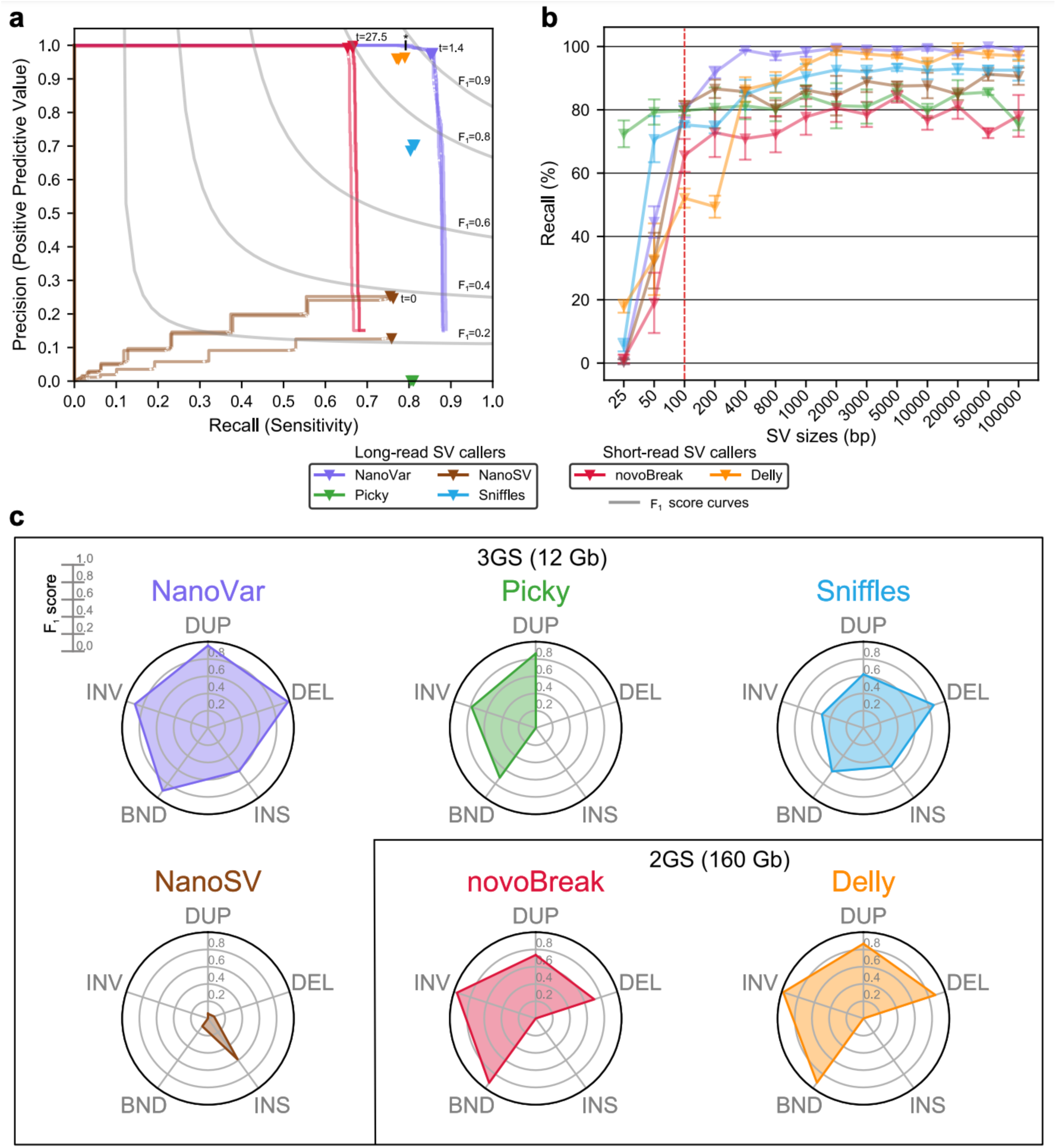
NanoVar performance benchmarking. (**a**) Precision and recall of SV detection by SV caller tools in simulation data (Three datasets with about 1150 SV each, sizes from 25 bp to 100k bp). The representation of curves and optimal confidence threshold score (labeled as t) are only depicted for tools which have confidence scoring for each breakend. The asterisk (*) on the NanoVar curve marks the confidence threshold score of t=2.6, which is the threshold used for higher stringency in patient data. (**b**) SV recall of varying SV sizes (in base pairs) detected by each tool in simulation data. Vertical red dotted line demarks SV of size 100 bp. (**c**) Radar charts showing the F_1_ scores for each SV class characterized by each tool for SV sizes 100 bp to 100k bp in simulation 1. DUP: tandem duplication, DEL: deletion, INS: insertion, BND: breakend, INV: inversion.

**Figure 3:**
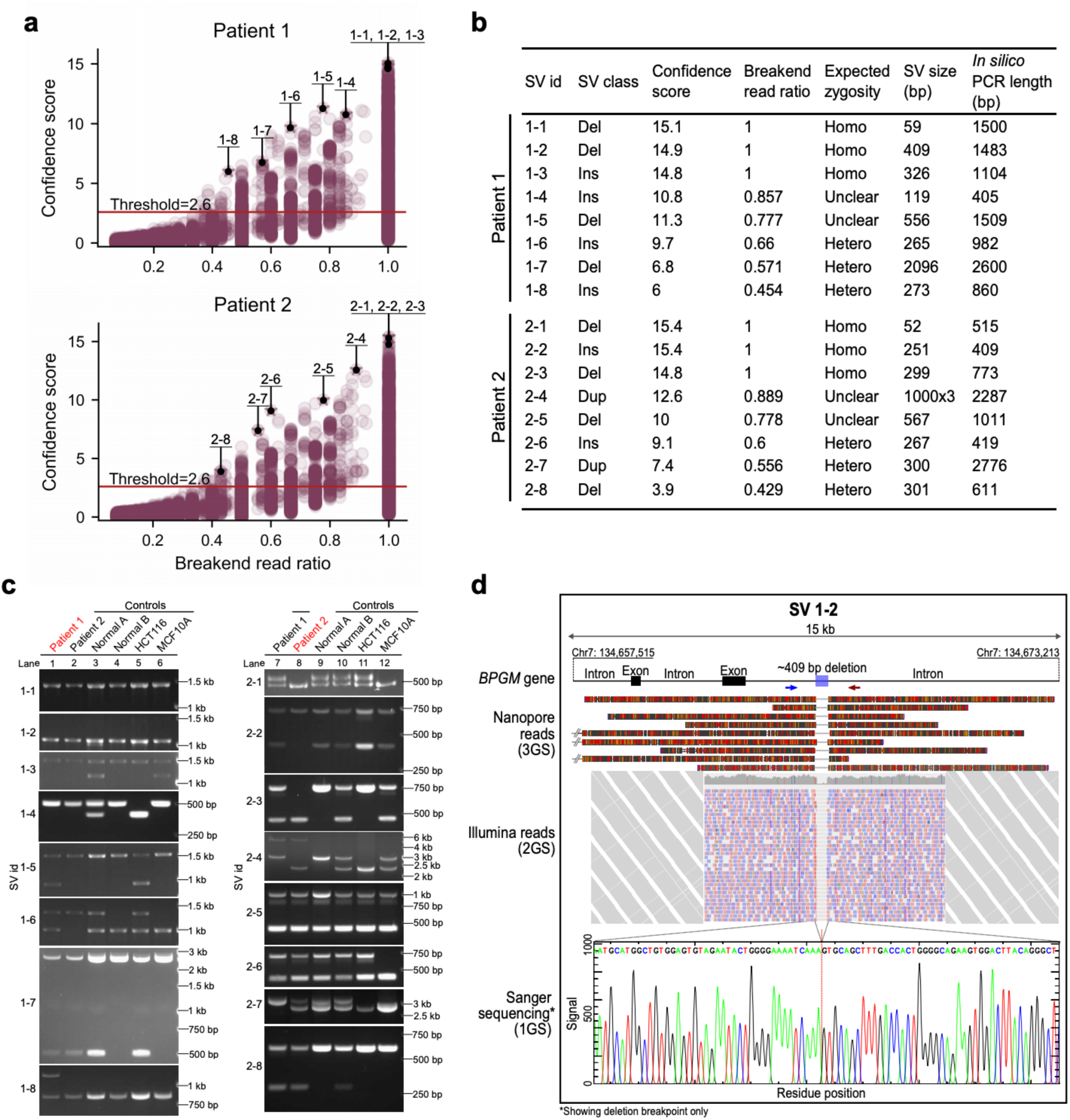
Precise patients’ SV characterization by NanoVar (and SV zygosity prediction.) (**a**) Scatter plots showing the confidence score and breakend read ratio of each SV characterized in Patient 1 (top) and Patient 2 (bottom). SVs selected for validations are labeled on the plots by their SV id. The red horizontal line indicates the confidence score threshold used for filtering. (**b**) Table displaying the details of SVs selected for validation for Patient 1 and Patient 2. (**c**) Gel electrophoresis images of PCR products corresponding to each of the SVs in table (b), amplified from the genomic DNA of Patient 1 and 2, normal donors (Normal 1 and Normal 2) and cell lines (HCT116 and MCF10A). Sample names in red (left image lane 1, right image lane 2) indicate the sample where the SV was initially detected. (**d**) Schematic illustrating a 409 bp deletion (SV 1-2) in the intronic region of the gene *BPGM* in Patient 1, supported by 3GS Nanopore reads (top), 2GS Illumina reads (middle) and 1GS Sanger sequencing chromatogram (bottom). Blue and red arrows represent the primer locations used for PCR amplification. For each Nanopore read, base substitutions and base insertions are represented by red and orange markers respectively. Base deletions are represented by gaps. All Nanopore reads have at least 90% alignment identity. Illumina paired-end short reads are represented by pink (forward) and blue (reverse) small rectangles and the read coverages are displayed in grey above all the reads. The red dotted line on the sequencing chromatogram marks the precise breakpoint of the deletion at single nucleotide

### Precise SV characterization in AML patients using NanoVar

We tested the NanoVar workflow on two Asian patients diagnosed with acute myeloid leukemia (AML) (Patient 1 and Patient 2) to evaluate the workflow’s feasibility and SV characterization accuracy in low sequencing depth clinical samples. Genomic DNA extracted from bone marrow mononuclear cells of each patient was sequenced by Nanopore sequencing using two to five MinION flowcells, generating about 12.4 gigabases (Gb) and 12.3 Gb of sequencing data respectively (Supplementary Table 3). The sequencing reads were applied to NanoVar for read mapping and SV characterization. A total of 7030 SVs (1794 DUPs, 2296 DELs, 2320 INSs, 493 BNDs, 127 INVs) were characterized in Patient 1, and 7761 SVs (1694 DUPs, 3361 DELs, 2098 INSs, 493 BNDs, 115 INVs) in Patient 2 (Supplementary Fig. 4). To evaluate NanoVar’s accuracy in these samples, we surveyed eight SVs from each patient for PCR validation. The SVs were selected according to their confidence score and breakend read ratio which estimates for SV zygosity. A single top confidence scoring SV was selected from each breakend read ratio interval of 0.1 from 0.4 to 0.9, while the top three highest confidence scoring SVs were selected for the interval of 0.9 to 1.0 (Fig. 3a). All the selected SVs are situated in autosomal chromosomes (Supplementary Table 4). Primer sequences were designed flanking the breakend location(s) of each SV according to the reference genome and their referenced amplicon lengths (*in silico* PCR length) are recorded in Figure 3b, along with their breakend read ratios and estimated SV sizes. PCR performed for each SV in their respective patient samples revealed that all 16 SVs were validated to be true based on their product size deviation (Fig. 3c lanes 1 and 8, Supplementary Table 5a). Moreover, the PCR results were agreeable with the SV class, SV size, and SV zygosities estimated by NanoVar, based on the number of PCR products (one product for homozygous, two products for heterozygous). All amplified products were gel extracted and their sequence identity validated by Sanger sequencing (Supplementary data). Besides PCR validation, all 16 SVs were also found to be supported by mapped 2GS short-reads generated by Illumina WGS (Supplementary Fig 5). Figure 3d illustrates an example of how a deletion SV (SV 1-2), characterized by 3GS Nanopore long-reads, can be supported by 2GS Illumina short-reads and 1GS Sanger sequencing. The successful validation of SVs with varying breakend read ratios and varying confidence scores affirms NanoVar’s precision in SV characterization in clinical samples.

### NanoVar recalls SVs discovered by PCR

We went on to uncover shared SVs between Patient 1 and Patient 2 amongst the selected SVs by testing the SVs reciprocally in each patient by PCR using the same primers and cycle conditions (Fig. 3c lanes 7 and 2). Patient 1 was discovered to possess all eight SVs that were validated in Patient 2, while Patient 2 possesses six of the eight SVs validated in Patient 1. In total, 14 out of 16 validated SVs were discovered to be shared between Patient 1 and Patient 2, leaving SV 1-5 and 1-8 to be Patient 1 specific (Fig. 3c, Supplementary Table 5). Next, we investigated if NanoVar was sensitive enough to capture these shared-SVs in the respective patient. Among the eight shared-SVs in Patient 1 and six shared-SVs in Patient 2, NanoVar was able to capture five shared-SVs of each, aggregating to 10 out of 14 shared-SVs detected (Supplementary Table 5). We investigated the four undetected shared-SVs and found out that three of them (SVs 2-2, 2-3, 1-7) were in fact captured by NanoVar but did not pass the confidence score threshold of 2.6, possibly due to insufficient read depth coverage (3X to 5X) for heterozygous SV detection. The other undetected heterozygous shared-SV (SV 2-8) was not detected due to the absence of SV-associated reads at its locus. This SV locus was found to be under-sequenced, covered only by two reads that likely originated from the wild-type allele in Patient 1.

### NanoVar characterizes polymorphic SVs

Upon assessing the presence of these SVs across more samples, we observed that most SVs appeared to be polymorphic. We tested the SVs in several samples consisting of normal hematopoietic stem cells (HSCs) from two non-AML Asian individuals (Normal A and Normal B), an epithelial colorectal carcinoma cell line (HCT116), and a Caucasian non-tumorigenic breast epithelial cell line (MCF10A). We found out that normal HSC samples do possessed many of the SVs (13 out of the 16 SVs) which existed in at least one allele in either one or both HSC samples, while SVs 1-5, 1-8, and 2-4 were absent in both HSC samples (Fig. 3c lanes 3, 4, 9, 10). Interestingly, these 13 shared-SVs were also present in both Patient 1 and Patient 2, suggesting that these SVs might be prevalently found in cells, irrespective of AML disease. On the contrary, SVs absent in the HSC samples (SVs 1-5 and 1-8) were exclusively found in Patient 1, except for SV 2-4, which was found in both patient samples. Out of these 13 shared-SVs, 11 of them were also present in non-hematopoietic cell lines such as HCT116 and MCF10A (Fig. 3c lanes 5, 6, 11, 12), leaving two SVs (SV 2-7, 2-8) undetected in either cell lines. Taken together, the majority of the SVs (11 out of 16 SVs) were commonly found across the samples regardless of AML disease status, cell type, and ethnicity. Moreover, most SVs exhibited different zygosities among the samples (All SVs excluding SV 1-1, 1-2, 1-8, 2-5), which portrayed the polymorphic nature of these SVs. Within the scope of this study, we categorized the SVs into three groups based on their prevalence and zygosity variation across the samples: (1) Rare sample-specific SVs (SV 1-8), (2) Common SVs with no zygosity variation (SV 1-1, 1-2, 2-5), and (3) Common SVs with zygosity variation or polymorphic SVs (remaining SVs) which constituted most of the SVs characterized by NanoVar.

### The NanoVar workflow is time efficient

We compared the CPU time and maximum resident set size (memory) used by the workflows of each tool for SV characterization in Patient 1 to evaluate their processing speed and memory usage (Supplementary Table 6). Among 3GS SV callers, NanoVar stood out as the most time efficient tool by requiring about 10-fold lesser CPU hours than the rest to process 12 Gb of sequencing data using 24 threads. In real time, NanoVar took 194 minutes for the entire analysis of Patient 1 which is the fastest amongst all other tools. In exchange for its speed, NanoVar employs about 1.7-fold more memory than the rest, having a higher memory cap of 32 gigabytes.

## Discussion

NanoVar is a novel SV characterization tool that excels in accuracy and speed while overcoming the low-depth and error-prone sequencing of 3GS WGS. We showed that NanoVar can achieve a high SV detection accuracy (Precision: 0.97, Recall, 0.85) when using only 4X coverage datasets in simulations, which was observed to outperform existing 3GS SV callers. However, like most SV callers, NanoVar fails to resolve many small SVs or indels shorter than 100 bp in length and would require future optimizations in algorithm and technological improvements in ONT basecalling error-rate. NanoVar’s performance in the simulation was also reflected in low-depth patient data where we successfully validated a small subset of SVs discovered by NanoVar (16/16) and showed that its estimations on SV class, size and zygosity were reliable.

One major advantage of 3GS over 2GS SV calling approaches is the amount of raw sequencing data consumed. In our study, we showed that 12 Gb of 3GS data (4X coverage) produced a more comprehensive SV detection outcome than 160 Gb of 2GS data (50X coverage) when comparing analysis done by NanoVar and 2GS SV callers (Fig. 2c). The considerable reduction in sequencing data requirements could speed up SV analysis and reduce computational resources. 3GS approaches may be used in large-scale SV-association studies or routine sequencing-based clinical investigations to analyze and store massive amounts of sequencing FASTQ/FASTA files more efficiently^5,7^.

Despite NanoVar’s high accuracy, many of its characterized SVs might be SV polymorphisms commonly found in the human population. We observed that most of the validated SVs found in our AML patients also existed with mixed zygosities in normal HSC samples and other cell lines, suggesting that they might be benign polymorphic SVs. As SV polymorphisms are widespread in the human genome^27–32^, it is important to annotate these SVs by cross-referencing to collective polymorphic-SV databases to facilitate the discovery of disease-associated SVs. Alternatively, the GRCH38 human reference genome could be improved to encompass polymorphic sequence variations where polymorphic SVs could be readily identified^33^. The use of low-depth Nanopore sequencing for accurate and routine SV characterization could supply a steady flow of knowledge to the construction of such cohort reference genome and inclusive SV databases.

## Material and methods

### The NanoVar pipeline

NanoVar takes as input a WGS long-read FASTQ/FASTA file (at least 12 gigabases) and a reference genome and outputs two variant calling format (VCF) files (Total SV and filtered SV) and an HTML summary report. The NanoVar workflow comprises of three main stages: 1) long-read sequence mapping, 2) SV characterization with read-depth calculation and 3) artificial neural network (ANN) inferencing from a simulation-trained model.

#### Stage 1: Long-read sequence mapping

The first stage aligns long-read sequences to a user-provided reference genome using the tool HS-BLASTN^22^ (version 0.0.5+). HS-BLASTN is an accelerated sequence alignment search tool that uses the MegaBLAST algorithm. We selected HS-BLASTN over other long-read aligner tools because of its faster computational speed and accurate read alignment, based on our evaluation. Before running HS-BLASTN, tools from NCBI-BLAST+ are used to build a blast database (makeblastdb^34^, version 2.6.0+) and mask highly repetitive sequences (windowmasker^34,35^, version 2.6.0+). HS-BLASTN is run with the following parameters: “-reward 2 -penalty -3 -gapopen 0 - gapextend 4 -max_target_seqs 3 -outfmt 6”. The output is a BLAST-like tabular file containing alignment information of each read. Due to overlapping alignments within some reads, a Python script is used to trim the overlapped regions or select the best alignment based on alignment bitscore.

#### Stage 2: SV characterization and read-depth calculation

The alignment anchor sequences and divergent sequences/gaps of each read are analyzed by Python scripts to detect reads containing *novel adjacencies* (reads possessing split-read or hard-clipped alignments) and subsequently characterizes their SV class. A *novel adjacency* is defined as two adjacent genomic coordinates in a sample genome that are not found to be adjacent in the reference genome. A novel adjacency is represented as two genomic coordinates in the reference genome, each known as a *breakend*. We use an algorithm of conditional control statements for novel adjacency detection and SV characterization, described in Supplementary Figure 1. Any read that is found to possess a novel adjacency is labeled as an SV-associated read, otherwise, labeled as a normal read. Next, the read-depth was calculated at each breakend for SV-associated reads and normal reads separately. This gives us the number of breakend-supporting reads *B* and breakend-opposing reads *O* at each breakend. Due to repetitive sequences in the genome, artificial breakends with unusually high *B* may be falsely detected. In order to filter-out these untrue breakends, we define the upper limit of *B* as *U*, where breakends with *B* > *U* are considered outliers and removed. *U* is calculated by

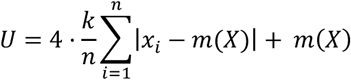

where *n* is the total number of genomic locations chosen, *x*_*i*,_ is the read-depth at genomic location *i, m*(*X*) is the median read-depth of all chosen genomic locations, and *k* is the constant scale factor 1.4826. The value of *U* is defined as four times the mean absolute deviation around the median (MAD) from the median in the distribution of a breakend read-depth assessment. This outlier detection method is an adaptation from Leys *et al*. where they proposed that the median absolute deviation is a more robust measure of dispersion than the standard deviation^36^. In our method, we use the MAD instead of the median absolution deviation to reduce fluctuations caused by discontinuous median integers. The breakend read-depth assessment is a sampling procedure to approximate the read-depth of SV throughout the genome. It is performed by randomly choosing *n* number of genomic locations and calculating the number of reads covering each location after adjusting for *G*. This produces a distribution similar to a gamma distribution and the median *m*(*X*) and MAD can be computed. According to our simulations, we empirically defined *U*, the deviation of more than four times the MAD from the median *m*(*X*), to be an outlier threshold, in the context of the human genome. Hence, any breakend which has *B* greater than *U* will be omitted and the remaining breakends will proceed to the next stage of ANN inferencing.

#### Stage 3: ANN inference

A trained ANN model is employed to improve SV characterization accuracy by evaluating read alignment characteristics and breakend read-depth information. For each novel adjacency, 23 scaled features are inferred by the ANN model which produces an inference value *P* ranging from 0 to 1. Next, *P* is exponentially scaled inversely according to the value of *B* and the final predicted score *S* is expressed logarithmically related to its error rate. S is described as

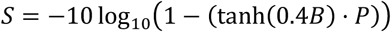

where *B* is the number of breakend-supporting reads at a novel adjacency and *P* is the ANN inference value of a novel adjacency. The hyperbolic tangent function is used to decrease the value of *P* non-linearly when *B* is low (*B* = [1, 2, 3]), as a low *B* confers low confidence. The value of *S* is proportional to the confidence level of a novel adjacency and is used to filter confident novel adjacencies from the total VCF output file to create the filtered VCF output file. A HTML summary report is also generated at the end of each run.

### Artificial neural network model and training

The features used by the ANN are described below (number in parentheses represent the number of neurons):

- Aligned/unaligned percentages flanking the novel adjacency (5)
- Alignment E-values flanking the novel adjacency (2)
- Relative alignment bit scores flanking the novel adjacency (2)
- Alignment identities flanking the novel adjacency (2)
- The fraction of mismatches in alignments flanking the novel adjacency (2)
- A fraction of gaps in alignments flanking the novel adjacency (2)
- SV complexity -number of coexisting SV found at the novel adjacency (1)
- Total number of alignments found on read (1)
- Total number of SV that seemed to be captured by read (1)
- Number of different chromosomes the read aligns (1)
- The fraction of alignments less than 5% of read length (1)
- Number of breakend-supporting reads *B* (1)
- A fraction of breakend-supporting reads *B* over total read depth *B*+*O* (1)
- If SV is an insertion/deletion, the size of the inserted/deleted segment (1)

The value of each feature is scaled to the range of [0, 1] by min-max normalization. The Python library Keras^37^ was used to build and infer the ANN model. The backend engine used with Keras is TensorFlow^38^. The neural network model is a feed-forward network consisting of a 23 neurons input layer, two hidden layers of 12 and 5 neurons sequentially, and a single neuron output layer. The Rectified Linear Unit (ReLU) activation function is used for the two hidden layers, while the Sigmoid activation function is used for the output layer. Dropout regularizations were implemented after each hidden layer with probabilities of 0.4 and 0.3 sequentially. If *y*_*K,i*_ denotes the value of the i-th neuron in the k-layer, we have that

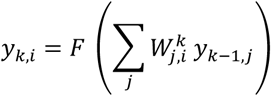

where F(x) = max(x,0) denotes the ReLU non-linearity and 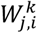 is the neural weight between the j-th neuron of the (k-1)-th layer and the i-th neuron of the k-th layer.

*In silico* 3GS reads from a simulated genome consisting of 61,316 mixed zygosity SV was used to train a binary classifier ANN model through supervised learning. The training dataset consist of 933,124 true and 62,902 false examples of novel adjacencies. Another simulated dataset with a different SV profile was used as the test dataset. Binary cross entropy was used as the loss function and stochastic gradient descent (SGD) was used as the optimizer algorithm with their default parameters. The classification accuracy is collected and reported as the metric to assess the performance of the model. One hundred epochs were performed for the model training, with each epoch having 12,000 true and 12,000 false randomly selected examples and a batch size of 400 examples per iteration.

### SV genome simulation for test datasets

The template genome used for genome simulation consisted of the main nuclear chromosomes (Chromosome 1 to Y) in the GRCh38 human reference genome assembly with their gap regions (N regions) replaced by tandemly repeated sequences. Telomeric (TTAGGG)n^39^ sequence replaced the gap regions at chromosome ends, and consensus centromeric alpha DNA^40,41^ replaced the centromeric and remaining gap regions. The removal of gap regions will allow uniformity in read simulation and a proper simulation of a real sequencing library. The R Bioconductor package, RSVSim^42^, was used to introduce novel adjacencies systematically in a reference genome to create different classes of SV. Five classes of SV were introduced into the template genome at varying amounts: 650 deletions, 200 inversions, 100 tandem duplications with single duplication each, 150 human genomic sequence insertions and 50 viral sequence insertions (serving as novel sequence insertions). Viral sequences used for viral insertions were part of 54 viral genomes taken from GenBank^43^ (Supplementary Table 7). The virus selection was based on their ability to integrate into the host genome. The amounts for each SV were based approximately on SV occurrence statistics of clinical cancer genomes by Hillmer et al.^44^. To simulate SV sequence variability, each novel adjacency has a 20 bp flanking region where bases had a 25% chance of single nucleotide polymorphism (SNP) and a 50% chance of introducing indels with a maximum indel length of 5 bp. The location of each SV was randomly generated throughout the whole genome by *RSVSim*. The sizes and quantity for each class of SV are recorded in Supplementary Figure 2a. A total of three genomes were simulated and their FASTA files can be downloaded from doi:10.5281/zenodo.2599376 or http://dx.doi.org/10.5281/zenodo.2599376.

### Mix zygosity SV genome simulation for the training dataset

The mix zygosity SV genome was created by three simulated genomes with varying number of SV from the same SV profile: Genome A has 61,316 SV (100%), Genome B has 51,099 SV (83%), and Genome C has 30,659 SV (50%). The SVs in Genome C are a subset of SVs in Genome B. Different number of *in silico* 3GS reads were generated for each genome: 5 million reads from Genome A, 5 million reads from Genome B, and 10 million reads from Genome C. The combination of all the reads produced the simulation of homozygous SV (50%), heterozygous SV (33%), and low-confidence SV (17%). A homozygous SV only has breakend-supporting reads at their breakends while a heterozygous SV has both breakend-supporting and breakend-opposing reads at similar proportions. A low-confidence SV simulates a false SV event and has a majority of its breakend reads being breakend-opposing.

### *In silico* Third-Generation sequencing (3GS)

*Nanosim*^45^ was used to generate *in silico* 3GS reads from the simulated SV genomes. Read features, such as read length, SNP, and indel profile, were modeled according to that of real ONT MinION reads from Patient 1 and Patient 2, which are provided as input into *Nanosim*. Two million reads were generated for each SV genome. Comparison for read length and indel proportion between real reads and *in silico* generated reads are shown in Supplementary Figures 2b and 2c. Statistics of reads and genome mapping can be found in Supplementary Table 8.

### *In silico* Second-Generation sequencing (2GS)

DWGSIM^46^ was used to generate *in silico* 2GS reads from the three simulated SV genomes. The generation of 2GS reads followed these settings: Illumina platform, 307 bp average insert size, 59 bp standard deviation of insert size, 150 bp read length, paired-end reads, 50X mean coverage across all regions, uniformly increasing per base error rate from 0.1% at start of read to 1% at end of read, and contains no mutations, indels, or random DNA reads. The insert size, read length and coverage follow that of real whole genome 2GS data of Patient 1 and Patient 2.

Statistics of reads and genome mapping can be found in Supplementary Table 8.

### Performance evaluation in simulation datasets

For each simulation dataset, a filtered list of ground truth SV genomic coordinates (BED file with ± 400 bp window about each breakend coordinate) was used to evaluate SV detection precision and recall for each tool. The original ground truth SV list was filtered to remove SVs that were not covered by any long-read or SVs which fell into genomic gap regions due to random generation. For the output SVs from each tool, we removed any SV entry which corresponds to translocation/insertion SVs at genome gap junctions indirectly introduced during N region replacement for genome simulation. The intersection between ground truth and query SV coordinates were carried out by BEDTools^47^. Precision and recall were computed by Scikit-learn^48^ and F_1_ score was calculated by the equation:

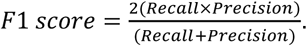

### DNA sample source

DNA samples used in this study were acquired from four individuals: two patients with AML (Patient 1, Patient 2) and two healthy donors (Normal A, Normal B). Informed consent from all subjects was obtained for genetic profiling such as whole genomic DNA sequencing. Patient 1 and Patient 2 had the M5 AML classification (Acute monocytic leukemia) with FLT-3 Asp835 mutations, but the absence of recurrent SV based on karyotyping. Patient 1 also has a mutation in the NPM1 gene. All subjects are of Asian ethnicity.

### Cell lines

The HCT116 and MCF10A cell lines were obtained from Horizon Discovery (HD PAR-007) and ATCC (ATCC CRL-10317™) respectively and grown in their respective recommended growth culture conditions.

### Genomic DNA extraction

Mononuclear cells (MNCs) of all individuals were isolated from bone marrow. Bone marrow from the pelvic bone was used for Patient 1 and 2, and bone marrow from the femur was used for Normal A and B. For Patient 1 and 2, bone marrow was diluted in phosphate-buffered saline (PBS) containing 2% HyClone™ Fetal Bovine Serum (FBS) (GE Healthcare Life Sciences) and 2 mM EDTA. MNCs were then isolated by Ficoll-Paque layering using Ficoll-Paque PLUS (GE Healthcare Life Sciences) following the manufacturer’s protocol. For Normal A and B, additional processing steps were carried out due to the presence of liquid fats. Femoral marrow was diluted in PBS containing 10% FBS, 3 mM EDTA and 0.4% sodium citrate. Cells were strained using a 100 µm cell strainer and pelleted by centrifugation at 300 g for 10 min at room temperature (RT) without acceleration and brakes. Red blood cells were lysed in 40 ml ACK lysis buffer (0.15 M NH_4_Cl, 1 mM KHCO_3_, 0.1 mM EDTA-Na_2_, pH adjusted to 7.2 - 7.4) at RT for 5 min. Cells were pelleted by centrifugation again with the same settings. The cell pellet was resuspended in PBS containing 2% FBS and 2 mM EDTA, and subsequently MNC isolation by Ficoll-Paque layering following the manufacturer’s protocol. MNCs of Normal A and B were enriched for hematopoietic stem cells (HSCs) by CD34 cell surface marker selection using the CD34 MicroBead kit, human (Miltenyi Biotec) according to manufacturer’s instructions. The buffer used for CD34+ cell selection is PBS containing 2% FBS and 2 mM EDTA. Genomic DNA of MNCs and CD34+ cells were extracted using AllPrep DNA/RNA/miRNA universal kit (Qiagen) and genomic DNA of HCT116 and MCF10A cells were extracted using conNorventional phenol-chloroform extraction method.

### Nanopore whole-genome sequencing and basecalling

High molecular weight genomic DNA (1-1.5 µg) was sheared to 6-10 kb fragments by the G-tube (Covaris). Library preparation was performed using ONT 1D or 2D Ligation Sequencing kits (SQK-LSK108, SQK-LSK208) following their protocol. FFPE DNA repair was not carried out. DNA ends were prepared using NEBNext Ultra II End Repair/dA-Tailing Module (New England Biolabs) for extended incubation time (30 min - 20°C, 30 min - 65°C). Ligation of sequencing adapters was performed using Blunt/TA Ligase Master Mix (New England Biolabs). Libraries were sequenced using the MinION sequencer on either R9.4 or R9.5 flowcells for 48 h without local base-calling. Base-calling was carried out by Metrichor or Albacore. Details of sequencing runs are documented in Supplementary Table 3. FASTQ/FASTA files were extracted from FAST5 files using h5dump (version 1.8.16) from HDF5 tools^49^. For the 2D protocol, the FASTQ/FASTA was extracted from the template strand instead of the combined strand if the complementary strand failed in quality.

### Nanopore read mapping and SV calling

For SV calling with NanoSV^24^ (version 1.1.6), LAST (version 938) was used to map reads to the reference genome with default parameters using 24 threads. The scoring parameters for LAST were generated from a 20,000 reads subsample using last-train. NanoSV was run with the default configuration parameters using 24 threads and we input our own hg38 random BED file for coverage depth calculations. We obtained the confidence score for each breakend from their QUAL value found in the output VCF file. We called SV with Picky^23^ (version 0.2.a) using the BASH script they provided with 24 threads. We used their recommended LAST parameters for read mapping: “-C2 -K2 -r1 -q3 - a2 -b1 -v -v”. Picky was run with default parameters as in the BASH script. For SV calling with Sniffles^19^ (version 1.0.8), NGMLR (version 0.2.6) was used for read mapping with default parameters and 24 threads. Sniffles was run using 24 threads with the -s 2 parameter which allowed at least two reads as minimum support for an SV to be reported. All SAM file sorting, BAM conversion and BAM indexing were carried out by SAMtools^50^. For calculating read mapping statistics, BWA-0.7.15^51^ was used for read alignment with the BWA-MEM parameter “-x ont2d” and statistics were calculated using SAMTools^50^.

### Illumina whole-genome sequencing, mapping and SV calling

Genomic DNA (1 µg) was randomly sheared to 350 bp fragments with Covaris cracker (Covaris) followed by sequencing library preparation using the Truseq Nano DNA HT Library Prep kit (Illumina). Sequencing libraries were sequenced paired-end 150 bp on the HiSeq X Ten sequencing platform (Illumina) with the HiSeq X Ten Reagent Kit v2.5 (Illumina) to a mean depth of coverage of about 50x. Reads were mapped to GRCh38 genome assembly using BWA-0.7.17^51^ with the default BWA-MEM parameters and 24 threads. SAM files were processed to sorted and indexed BAM files using SAMtools^50^. For SV calling with novoBreak^25^ (version 1.1.3rc), sorted and indexed BAM files were input with default run parameters using 24 threads. A dummy BAM file was simulated (GRCh38) to be used as a matched normal control. The confidence score for each breakend was obtained from the QUAL scores in the output VCF file. For SV calling with Delly^26^ (version 0.7.8), duplicated reads in the BAM files were identified by Picard MarkDuplicates^52^ before running Delly with the provided hg38 exclude file and its default parameters.

### SV experimental validation

Polymerase chain reaction (PCR) was carried out to amplify SV-containing regions in the genomes of each sample. We used two different PCR master mixes. REDiant 2X PCR Master Mix (Axil Scientific) was used for conventional PCR amplification, whereas LongAmp Taq 2X Master Mix (New England Biolabs) was used for longer (>1.5 kbp) or AT-rich PCR products. DMSO was added to a final concentration of 3% to increase the success rate of GC-rich product amplification. Primer sequences were designed using PrimerQuest Tool by Integrated DNA Technologies and shown in Supplementary Table 9. Forward and reverse primers were added to a final concentration of 0.4 µM each. 2 - 5 ng of genomic DNA was used as the template in each 25 µl PCR reaction. Standard three-step PCR settings were used for most PCR reactions on a thermal cycler. Touchdown PCR conditions may be implemented for some reactions to reduce unspecific products. PCR products were separated on 1% agarose TBE ethidium bromide gel by gel electrophoresis and DNA bands were visualized by UV light. DNA fragments were excised and extracted using a cotton wool gel filtration protocol as described in Sun et al. 2012 or QIAquick Gel Extraction Kit (Qiagen). DNA was subsequently purified using Agencourt AMPure XP beads (Beckman Coulter) following their protocol for PCR purification. Primary or nested PCR product sequences were validated by Sanger sequencing.

### CPU time and maximum memory consumption assessment

GNU Time (version 1.7) was used to assess the CPU time and maximum memory consumption of each tool. We assessed each tool by executing the following command: ‘/usr/bin/time –verbose – output=output.txt sh -c “Tool command”‘, and the results are stored in the output.txt file. The CPU time is calculated by combining the user and system time, and the maximum resident set size is taken as the maximum memory consumption.

## Code availability

NanoVar is an open-source free software available at GitHub (https://github.com/benoukraflab/NanoVar), licensed under the GNU Public License.

## Acknowledgements

Work in the T.B. laboratory is supported by the National Research Foundation, the Singapore Ministry of Education under its Centres of Excellence initiative and the RNA Biology Center at the Cancer Science Institute of Singapore, NUS, as part of funding under the Singapore Ministry of Education’s AcRF Tier 3 grants [MOE2014-T3-1-006]. This research was undertaken, in part, thanks to funding from the Canada Research Chairs program. C.Y.T. and R.T.M. are supported by a Doctoral Scholarship from the Cancer Science Institute of Singapore.

## Author contributions

C.Y.T. and T.B. designed the package. C.Y.T. programmed the NanoVar package with R.T.M.’s contribution. C.Y.T. performed all Nanopore sequencing data generation and analysis. C.Y.T., D.G.T. and T.B. interpreted the data. A.T. contributed to the machine learning module establishment. Y.G., M.J.F., B.T.H.K., W.W., C.H.N. and W.J.C. were involved in patients’ sample collection, DNA extraction and whole genome sequencing (Illumina technology). C.Y.T. and T.B wrote the manuscript. T.B. directed the project.

## References

1. Hurles, M. E., Dermitzakis, E. T. & Tyler-Smith, C. The functional impact of structural variation in humans. Trends Genet. 24, 238–45 (2008).

2. Weischenfeldt, J., Symmons, O., Spitz, F. & Korbel, J. O. Phenotypic impact of genomic structural variation: insights from and for human disease. Nat. Rev. Genet. 14, 125–138 (2013).

3. Redon, R. et al. Global variation in copy number in the human genome. Nature 444, 444–454 (2006).

4. Sudmant, P. H. et al. An integrated map of structural variation in 2,504 human genomes. Nature 526, 75–81 (2015).

5. Alkan, C., Coe, B. P. & Eichler, E. E. Genome structural variation discovery and genotyping. Nat. Rev. Genet. 12, 363–376 (2011).

6. Mitelman, F., Johansson, B. & Mertens, F. The impact of translocations and gene fusions on cancer causation. Nat. Rev. Cancer 7, 233–245 (2007).

7. Macintyre, G., Ylstra, B., Brenton, J. D. & Brenton, J. D. Sequencing Structural Variants in Cancer for Precision Therapeutics. Trends Genet. 32, 530–542 (2016).

8. Sanchis-Juan, A. et al. Complex structural variants in Mendelian disorders: identification and breakpoint resolution using short-and long-read genome sequencing. Genome Med. 10, 95 (2018).

9. Merker, J. D. et al. Long-read genome sequencing identifies causal structural variation in a Mendelian disease. Genet. Med. 20, 159–163 (2018).

10. Brandler, W. M. et al. Paternally inherited cis-regulatory structural variants are associated with autism. Science. 360, 327–331 (2018).

11. Miao, H. et al. Long-read sequencing identified a causal structural variant in an exome-negative case and enabled preimplantation genetic diagnosis. Hereditas 155, 32 (2018).

12. Andersen, C. L. et al. Frequent genomic loss at chr16p13.2 is associated with poor prognosis in colorectal cancer. Int. J. Cancer 129, 1848–1858 (2011).

13. Wang, Z.-Y. & Chen, Z. Acute promyelocytic leukemia: from highly fatal to highly curable. Blood 111, 2505–2515 (2008).

14. Goodwin, S., McPherson, J. D. & McCombie, W. R. Coming of age: ten years of next-generation sequencing technologies. Nat. Rev. Genet. 17, 333–351 (2016).

15. Croville, G. et al. Rapid whole-genome based typing and surveillance of avipoxviruses using nanopore sequencing. J. Virol. Methods 261, 34–39 (2018).

16. Ebbert, M. T. W. et al. Long-read sequencing across the C9orf72 ‘GGGGCC’ repeat expansion: implications for clinical use and genetic discovery efforts in human disease. Mol. Neurodegener. 13, 46 (2018).

17. Pendleton, M. et al. Assembly and diploid architecture of an individual human genome via single-molecule technologies. Nat. Methods 12, 780–786 (2015).

18. Seo, J.-S. et al. De novo assembly and phasing of a Korean human genome. Nature 538, 243–247 (2016).

19. Sedlazeck, F. J. et al. Accurate detection of complex structural variations using single-molecule sequencing. Nat. Methods 15, 461–468 (2018).

20. Tattini, L., D’Aurizio, R. & Magi, A. Detection of Genomic Structural Variants from Next-Generation Sequencing Data. Front. Bioeng. Biotechnol. 3, 92 (2015).

21. Liu, Q., Zhang, P., Wang, D., Gu, W. & Wang, K. Interrogating the ‘unsequenceable’ genomic trinucleotide repeat disorders by long-read sequencing. Genome Med. 9, 65 (2017).

22. Chen, Y., Ye, W., Zhang, Y. & Xu, Y. High speed BLASTN: an accelerated MegaBLAST search tool. Nucleic Acids Res. 43, 7762–7768 (2015).

23. Gong, L. et al. Picky comprehensively detects high-resolution structural variants in nanopore long reads. Nat. Methods 15, 455–460 (2018).

24. Cretu Stancu, M. et al. Mapping and phasing of structural variation in patient genomes using nanopore sequencing. Nat. Commun. 8, 1326 (2017).

25. Chong, Z. et al. novoBreak: local assembly for breakpoint detection in cancer genomes. Nat. Methods 14, 65–67 (2016).

26. Rausch, T. et al. DELLY: structural variant discovery by integrated paired-end and split-read analysis. Bioinformatics 28, i333–i339 (2012).

27. Kidd, J. M. et al. Mapping and sequencing of structural variation from eight human genomes. Nature 453, 56–64 (2008).

28. Kidd, J. M. et al. Characterization of missing human genome sequences and copy-number polymorphic insertions. Nat. Methods 7, 365–71 (2010).

29. Bailey, J. A., Kidd, J. M. & Eichler, E. E. Human copy number polymorphic genes. Cytogenet. Genome Res. 123, 234–43 (2008).

30. Antonacci, F. et al. Characterization of six human disease-associated inversion polymorphisms. Hum. Mol. Genet. 18, 2555–2566 (2009).

31. Handsaker, R. E., Korn, J. M., Nemesh, J. & McCarroll, S. A. Discovery and genotyping of genome structural polymorphism by sequencing on a population scale. Nat. Genet. 43, 269–276 (2011).

32. Audano, P. A. et al. Characterizing the Major Structural Variant Alleles of the Human Genome. Cell 176, 663–675.e19 (2019).

33. Paten, B., Novak, A. M., Eizenga, J. M. & Garrison, E. Genome graphs and the evolution of genome inference. Genome Res. 27, 665–676 (2017).

34. Camacho, C. et al. BLAST+: architecture and applications. BMC Bioinformatics 10, 421 (2009).

35. Morgulis, A., Gertz, E. M., Schaffer, A. A. & Agarwala, R. WindowMasker: window-based masker for sequenced genomes. Bioinformatics 22, 134–141 (2006).

36. Leys, C., Ley, C., Klein, O., Bernard, P. & Licata, L. Detecting outliers: Do not use standard deviation around the mean, use absolute deviation around the median. J. Exp. Soc. Psychol. 49, 764–766 (2013).

37. Chollet, F. & others. Keras. (2015).

38. Abadi, M. et al. TensorFlow: Large-Scale Machine Learning on Heterogeneous Systems. (2015).

39. Moyzis, R. K. et al. A highly conserved repetitive DNA sequence, (TTAGGG)n, present at the telomeres of human chromosomes. Proc. Natl. Acad. Sci. U. S. A. 85, 6622–6 (1988).

40. Vissel, B. & Choo, K. H. Human alpha satellite DNA--consensus sequence and conserved regions. Nucleic Acids Res. 15, 6751–2 (1987).

41. Bao, W., Kojima, K. K. & Kohany, O. Repbase Update, a database of repetitive elements in eukaryotic genomes. Mob. DNA 6, 11 (2015).

42. Bartenhagen, C. & Dugas, M. RSVSim: an R/Bioconductor package for the simulation of structural variations. Bioinformatics 29, 1679–1681 (2013).

43. Benson, D. A. et al. GenBank. Nucleic Acids Res. 45, D37–D42 (2017).

44. Hillmer, A. M. et al. Comprehensive long-span paired-end-tag mapping reveals characteristic patterns of structural variations in epithelial cancer genomes. Genome Res. 21, 665–75 (2011).

45. Yang, C., Chu, J., Warren, R. L. & Birol, I. NanoSim: nanopore sequence read simulator based on statistical characterization. Gigascience 6, 1–6 (2017).

46. Homer, N. DWGSIM: Whole Genome Simulator for Next-Generation Sequencing. GitHub repository (2010).

47. Quinlan, A. R. & Hall, I. M. BEDTools: a flexible suite of utilities for comparing genomic features. Bioinformatics 26, 841–842 (2010).

48. Pedregosa, F. et al. Scikit-learn: Machine Learning in {P}ython. J. Mach. Learn. Res. 12, 2825–2830 (2011).

49. The HDF Group. Hierarchical Data Format, version 5.

50. Li, H. et al. The Sequence Alignment/Map format and SAMtools. Bioinformatics 25, 2078–2079 (2009).

51. Li, H. & Durbin, R. Fast and accurate short read alignment with Burrows-Wheeler transform. Bioinformatics 25, 1754–1760 (2009).

52. Picard toolkit. Broad Institute, GitHub repository (2018).

53. Sun, Y., Sriramajayam, K., Luo, D. & Liao, D. J. A quick, cost-free method of purification of DNA fragments from agarose gel. J. Cancer 3, 93–5 (2012).

