## Supplementary Figures and Tables for "NanoVar: Accurate Characterization of Patients’ Genomic Structural Variants Using Low-Depth Nanopore Sequencing"

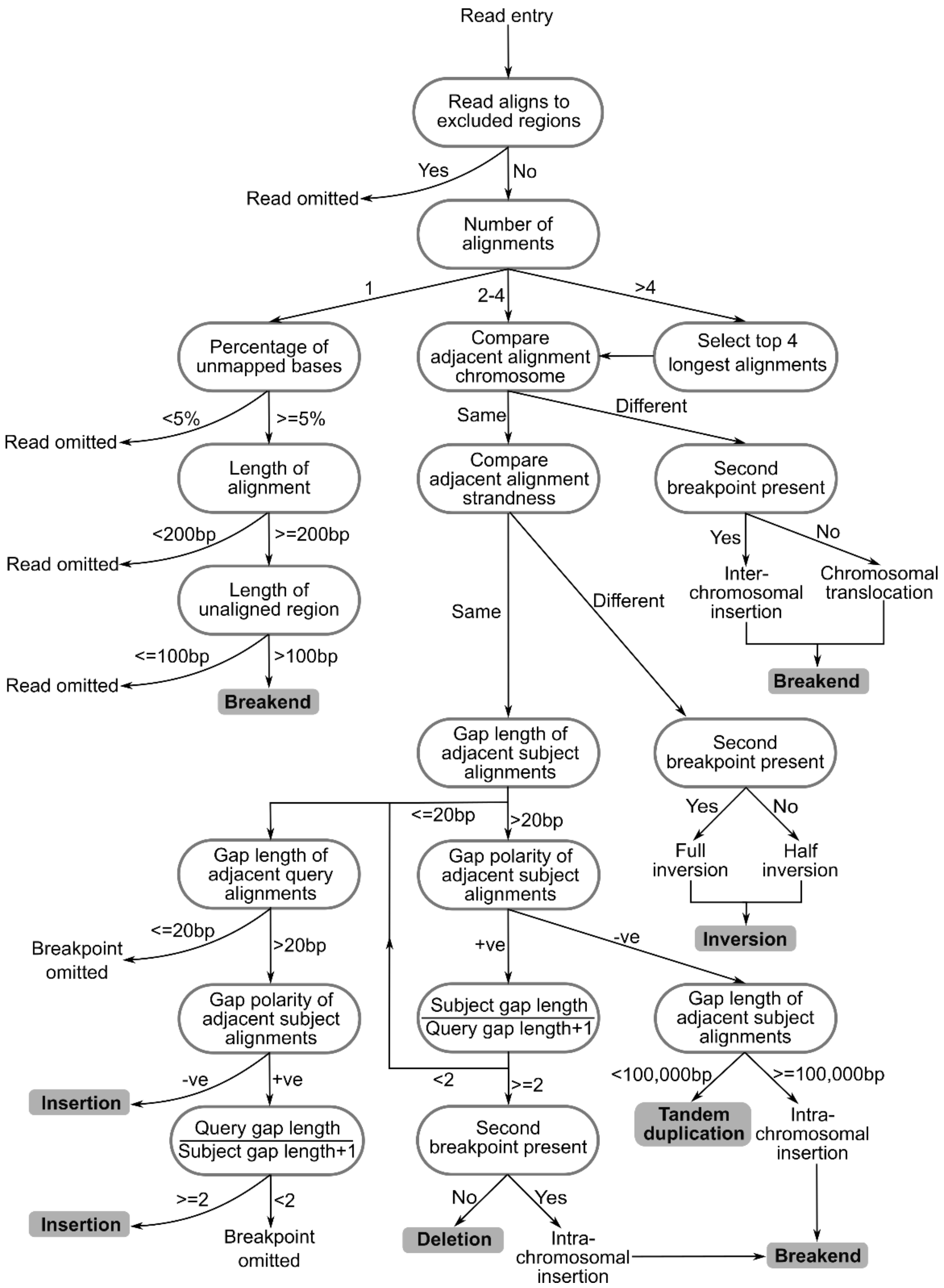


**Supplementary Figure 1**: NanoVar SV characterization algorithm displayed as a decision tree. The algorithm consists of only conditional control statements coded in Python to analysis long-read alignment profiles for SV characterization.

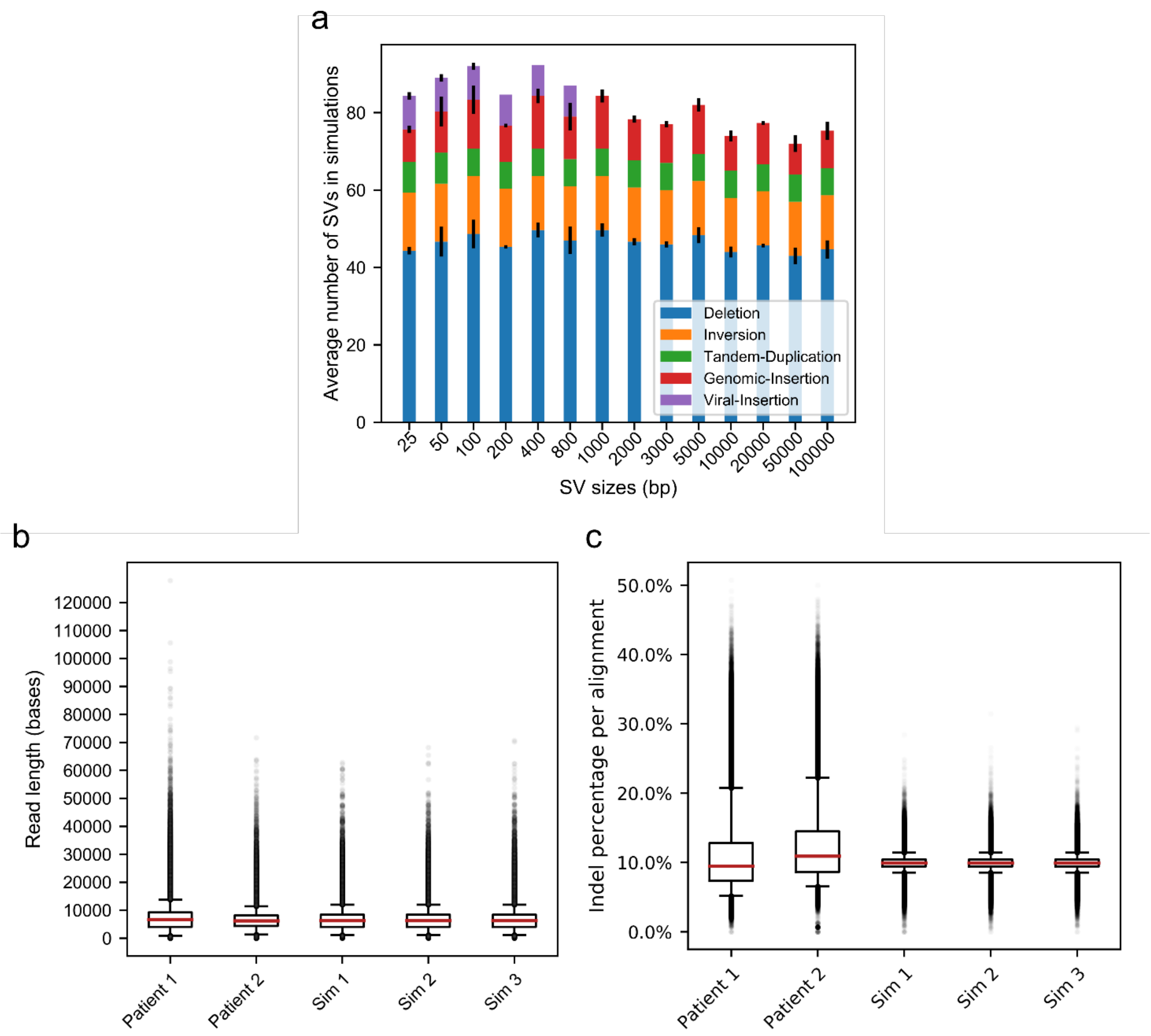


**Supplementary Figure 2:** SV simulation and read simulation information. **(a)** The average amount of SVs simulated in each dataset, separated by SV size and class. (**b)** Read length and **c)** indel percentage distribution of simulated long reads and whole-genome patient Nanopore sequencing reads.

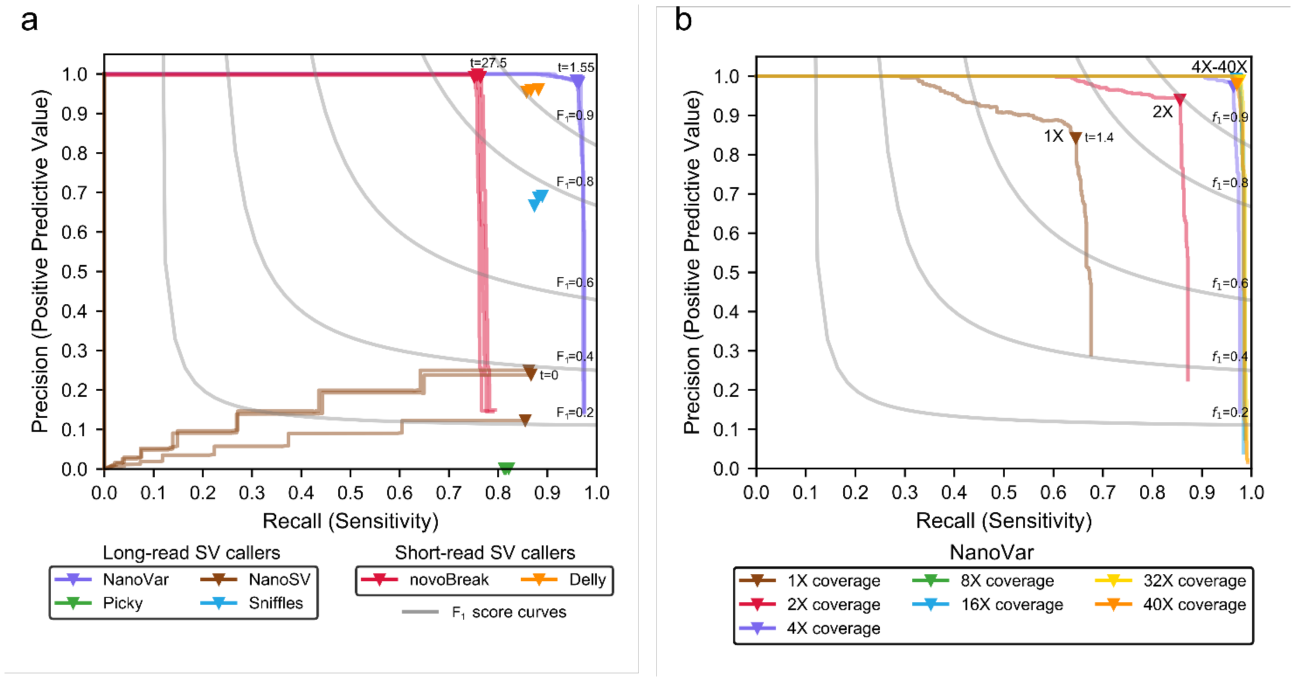


**Supplementary Figure 3**: **(a)** Precision and recall of different SV callers detecting simulated SVs with sizes equal to or more than 100 bp. NanoVar's recall improved greatly (from 85% to 96%) when small SVs (25-50 bp) were omitted. **(b)** Precision and recall of NanoVar's SV detection with different depths of coverage (1X to 40X) of input sequencing data. This was also carried out on simulated SVs with sizes equal to or more than 100 bp. All the triangular markers of each curve mark the consistent optimal confidence threshold score (t=1.4) across different coverages.

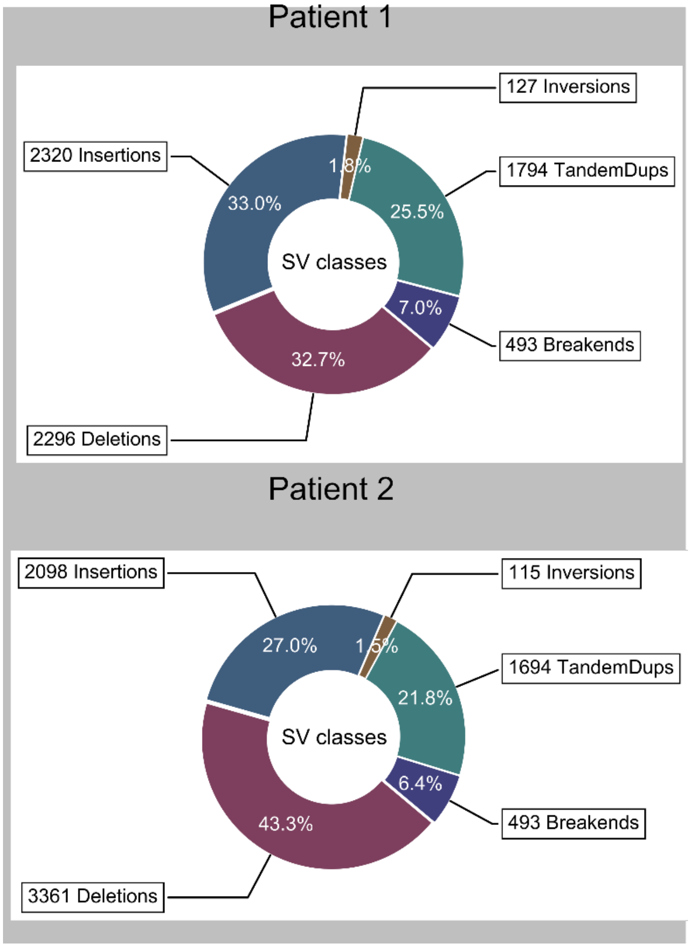


**Supplementary Figure 4**: Donut charts showing the distribution of SV classes characterized by NanoVar in Patient 1 and Patient 2.

a

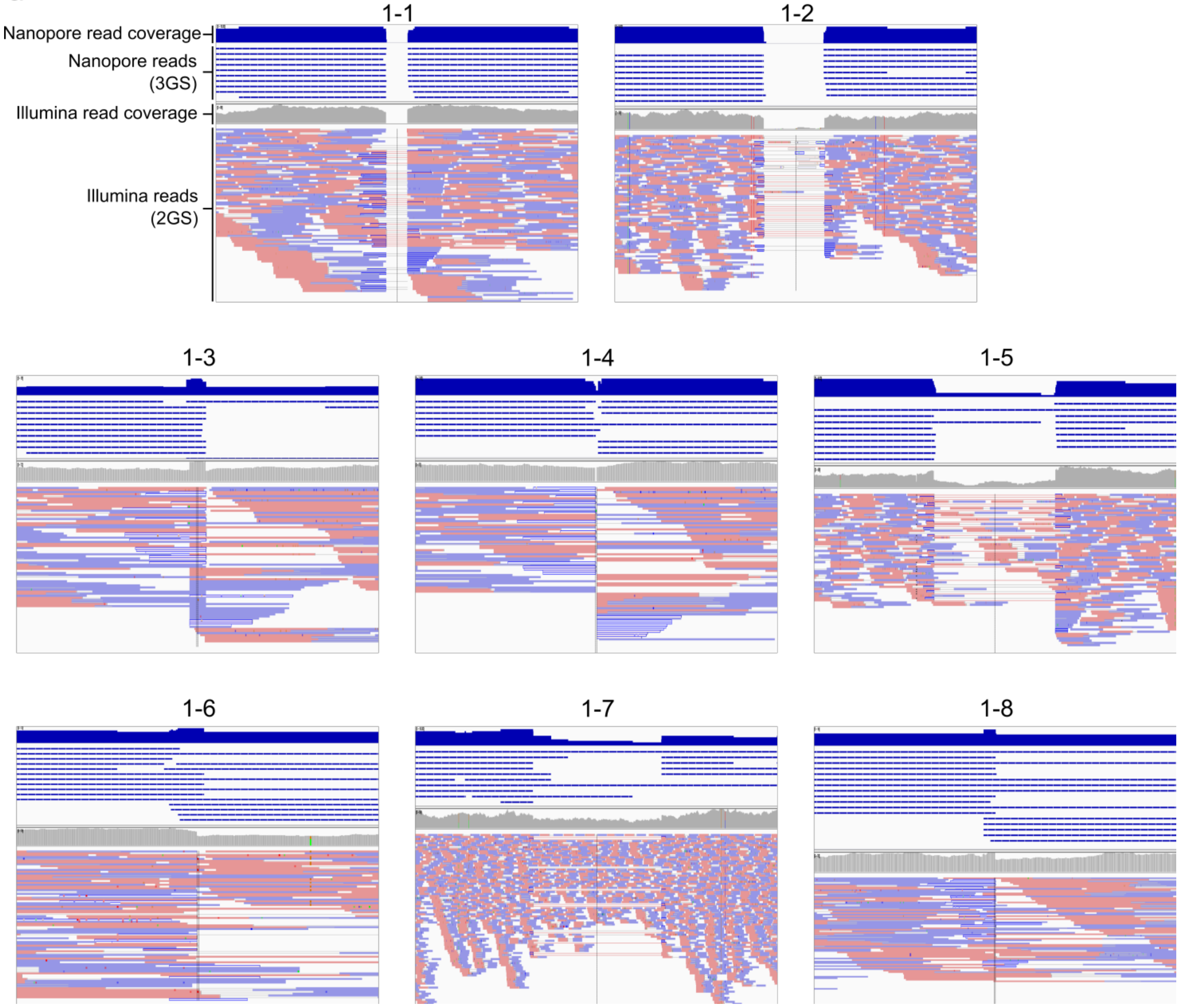
1-1
1-2
Nanopore read coverage
Nanopore reads
(3GS)
Illumina read coverage
Illumina reads
(2GS)
1-3
1-4
1-5
1-6
1-7
1-8

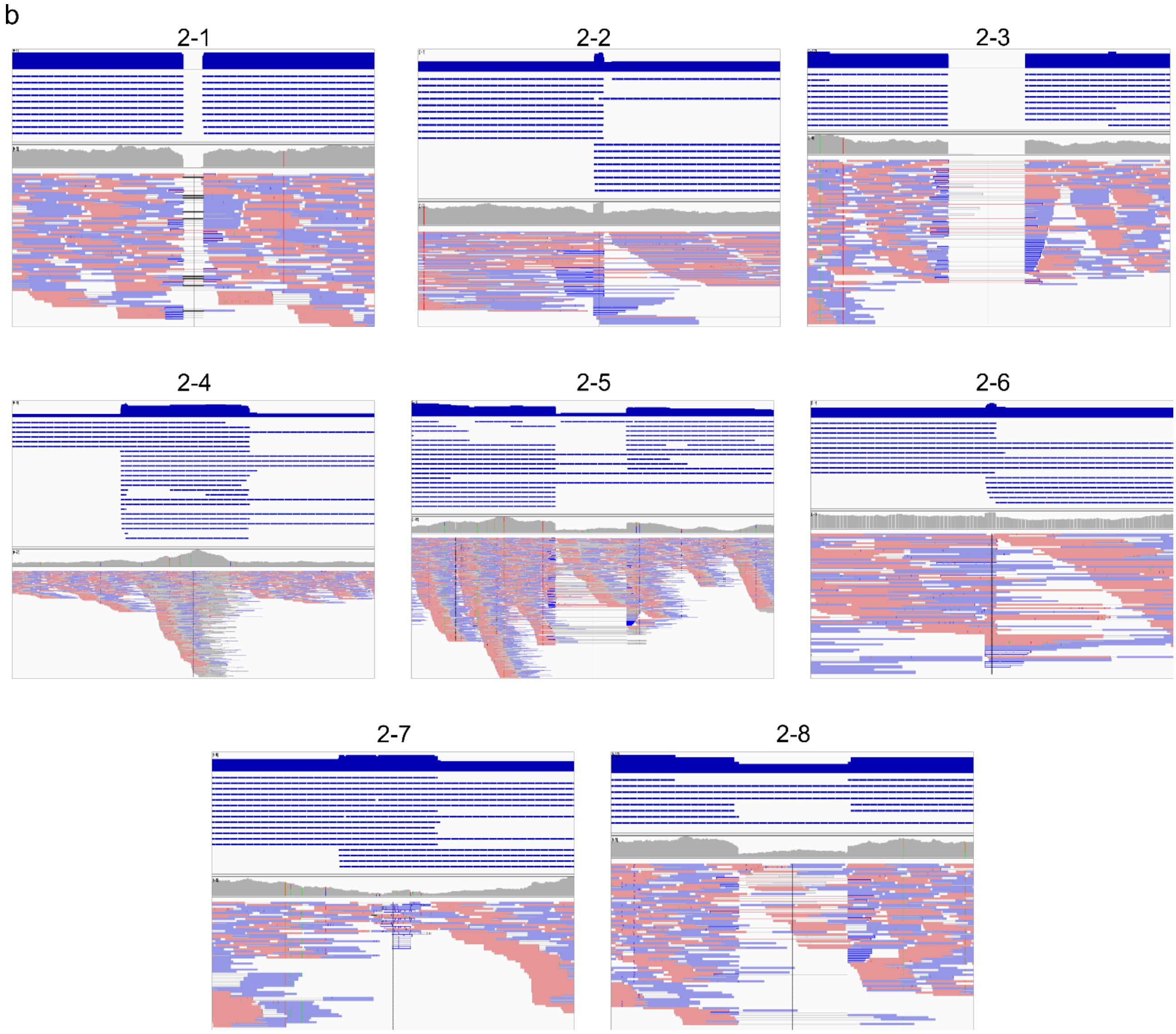


**Supplementary Figure 5**: Genome browser snapshots of Nanopore reads and Illumina reads aligning to hg38 reference genome at each SV breakpoint location for SVs in **(a)** Patient 1 and **(b)** Patient 2. The top half of each snapshot displays the Nanopore read alignments represented by blue rectangles while the bottom half displays Illumina reads represented by red and blue rectangles. The cumulative read coverages for Nanopore reads (blue) and Illumina reads (grey) are displayed above all the respective reads. The Integrative Genomics Viewer (IGV)[83,84] was used for the visualization of reads.

| Tool | Optimum threshold score | Recall | | | | Precision | | | | $F_1$ score | | | | Area under curve (AUC) | | | |
|---|---|---|---|---|---|---|---|---|---|---|---|---|---|---|---|---|---|
| | | Sim 1 | Sim 2 | Sim 3 | Avg | Sim 1 | Sim 2 | Sim 3 | Avg | Sim 1 | Sim 2 | Sim 3 | Avg | Sim 1 | Sim 2 | Sim 3 | Avg |
| NanoVar | 1.4 | 0.856 | 0.851 | 0.856 | 0.855 | 0.976 | 0.974 | 0.977 | 0.976 | 0.912 | 0.908 | 0.913 | 0.911 | 0.880 | 0.874 | 0.874 | 0.876 |
| NanoSV | 0 | 0.762 | 0.758 | 0.755 | 0.759 | 0.247 | 0.128 | 0.254 | 0.210 | 0.373 | 0.219 | 0.380 | 0.324 | 0.105 | 0.051 | 0.107 | 0.088 |
| Sniffles | N/A | 0.803 | 0.814 | 0.809 | 0.809 | 0.686 | 0.704 | 0.704 | 0.698 | 0.740 | 0.755 | 0.753 | 0.749 | N/A | N/A | N/A | N/A |
| Picky | N/A | 0.804 | 0.809 | 0.811 | 0.808 | 0.002 | 0.002 | 0.002 | 0.002 | 0.004 | 0.004 | 0.004 | 0.004 | N/A | N/A | N/A | N/A |
| novoBreak | 27.5 | 0.665 | 0.653 | 0.665 | 0.661 | 0.999 | 0.993 | 0.993 | 0.995 | 0.799 | 0.788 | 0.797 | 0.794 | 0.677 | 0.664 | 0.677 | 0.673 |
| Delly | N/A | 0.775 | 0.790 | 0.770 | 0.778 | 0.958 | 0.964 | 0.961 | 0.961 | 0.857 | 0.868 | 0.855 | 0.860 | N/A | N/A | N/A | N/A |

**Supplementary Table 1**: Precision and recall values of Figure 2a. Sim # refers to simulation dataset #.

| Tool | Optimum threshold score | DUP (Total :76) | | | | | DEL (Total: 497) | | | | | INS (Total: 32) | | | | |
|---|---|---|---|---|---|---|---|---|---|---|---|---|---|---|---|---|
| | | Recall | TP | TP+FP | Precision | $F_1$ score | Recall | TP | TP+FP | Precision | $F_1$ score | Recall | TP | TP+FP | Precision | $F_1$ score |
| NanoVar | 1.55 | 0.961 (73) | 76 | 80 | 0.950 | 0.955 | 0.984 (489) | 517 | 530 | 0.975 | 0.980 | 1.000 (32) | 34 | 76 | 0.447 | 0.618 |
| NanoSV | 0 | 0.803 (61) | 65 | 2321 | 0.028 | 0.054 | 0.038 (19) | 19 | 20 | 0.950 | 0.073 | 0.875 (28) | 33 | 75 | 0.440 | 0.586 |
| Sniffles | N/A | 0.763 (58) | 59 | 112 | 0.527 | 0.623 | 0.879 (437) | 439 | 520 | 0.844 | 0.861 | 0.938 (30) | 30 | 77 | 0.390 | 0.551 |
| Picky | N/A | 0.829 (63) | 63 | 70 | 0.900 | 0.863 | 0.781 (388) | 419 | 340420 | 0.001 | 0.002 | 0.813 (26) | 26 | 54911 | 0.000 | 0.001 |
| novoBreak | 27.5 | 0.632 (48) | 52 | 59 | 0.881 | 0.736 | 0.592 (294) | 342 | 374 | 0.914 | 0.719 | 0.000 (0) | 0 | 0 | 0.000 | 0.000 |
| Delly | N/A | 0.855 (65) | 65 | 74 | 0.878 | 0.867 | 0.847 (421) | 421 | 459 | 0.917 | 0.881 | 0.000 (0) | 0 | 0 | 0.000 | 0.000 |

| Tool | BND (Total : 115) | | | | | INV (Total: 159) | | | | |
|---|---|---|---|---|---|---|---|---|---|---|
| | Recall | TP | TP+FP | Precision | $F_1$ score | Recall | TP | TP+FP | Precision | $F_1$ score |
| NanoVar | 0.870 (100) | 203 | 218 | 0.931 | 0.900 | 0.824 (131) | 145 | 148 | 0.980 | 0.895 |
| NanoSV | 0.922 (106) | 203 | 3397 | 0.060 | 0.112 | 0.000 (0) | 0 | 0 | 0.000 | 0.000 |
| Sniffles | 0.670 (77) | 124 | 213 | 0.582 | 0.623 | 0.730 (116) | 116 | 298 | 0.389 | 0.508 |
| Picky | 0.565 (65) | 244 | 254 | 0.961 | 0.712 | 0.660 (105) | 105 | 109 | 0.963 | 0.783 |
| novoBreak | 0.930 (107) | 355 | 388 | 0.915 | 0.922 | 0.937 (149) | 343 | 345 | 0.994 | 0.965 |
| Delly | 0.870 (100) | 191 | 195 | 0.979 | 0.922 | 0.969 (154) | 305 | 306 | 0.997 | 0.983 |

**Supplementary Table 2**: Precision and recall values for different SV classes characterized by different tools presented in Figure 2c. DUP: tandem duplication, DEL: deletion, INS: insertion, BND: breakend, INV: inversion. Numbers in parentheses represent the number of true SVs recalled. SV class annotation accuracy is considered in this analysis.

| | Flowcell chemistry | ONT protocol | Basecaller | Amount of gDNA used | Number of flowcells | Number of reads | Total number of reads | Total bases ($x10^6$) |
|---|---|---|---|---|---|---|---|---|
| Patient 1 | R9.4 | 2D | Metrichor or Albacore | 7.3 µg | 2 | 441,637 | 1,793,667 | 12,407 |
| | R9.5 | 1D | Albacore | | 3 | 1,352,030 | | |
| Patient 2 | R9.5 | 1D | Albacore | 2 µg | 2 | 1,939,432 | 1,939,432 | 12,304 |

**Supplementary Table 3**: Oxford Nanopore MinION sequencing details of Patient 1 and Patient 2

| | SV id | SV class | Chromosome | Coordinates |
|---|---|---|---|---|
| Patient 1 | 1-1 | Del | 14 | 68528678-68528737 |
| | 1-2 | Del | 7 | 134665255-134665664 |
| | 1-3 | Ins | 18 | 40768187 |
| | 1-4 | Ins | 7 | 110844695 |
| | 1-5 | Del | 14 | 51911629-51912185 |
| | 1-6 | Ins | 14 | 53333449 |
| | 1-7 | Del | 4 | 79439473-79441569 |
| | 1-8 | Ins | 4 | 89221858 |
| Patient 2 | 2-1 | Del | 4 | 142108392-142108444 |
| | 2-2 | Ins | 4 | 137377851 |
| | 2-3 | Del | 5 | 79717888-79718187 |
| | 2-4 | Dup | 4 | 7834853-7835893 |
| | 2-5 | Del | 21 | 10475720-10476287 |
| | 2-6 | Ins | 10 | 35706138 |
| | 2-7 | Dup | 6 | 66974314-66974585 |
| | 2-8 | Del | 3 | 194894010-194894311 |

**Supplementary Table 4**: Genomic coordinates of SVs characterized in Patient 1 and Patient 2.

a

| | Patient 1 | | Patient 2 | |
|---|---|---|---|---|
| SV id | Validated | Shared-SV recalled | Validated | Shared-SV recalled |
| 1-1 | Y | n/a | n/a | Y |
| 1-2 | Y | n/a | n/a | Y |
| 1-3 | Y | n/a | n/a | Y |
| 1-4 | Y | n/a | n/a | Y |
| 1-5 | Y | n/a | n/a | Not shared |
| 1-6 | Y | n/a | n/a | Y |
| 1-7 | Y | n/a | n/a | N |
| 1-8 | Y | n/a | n/a | Not shared |
| 2-1 | n/a | Y | Y | n/a |
| 2-2 | n/a | N | Y | n/a |
| 2-3 | n/a | N | Y | n/a |
| 2-4 | n/a | Y | Y | n/a |
| 2-5 | n/a | Y | Y | n/a |
| 2-6 | n/a | Y | Y | n/a |
| 2-7 | n/a | Y | Y | n/a |
| 2-8 | n/a | N | Y | n/a |

b

| | SV validated | | Shared-SV recalled | |
|---|---|---|---|---|
| Sample | Individual | Total | Individual | Total |
| Patient 1 | 8/8 | 16/16 | 5/8 | 10/14 |
| Patient 2 | 8/8 | | 5/6 | |

**Supplementary Table 5:** SV validation in AML patients and NanoVar recall status for PCR-discovered SVs. **(a)** Results of SV validation and shared-SV detectability by NanoVar in the respective patient samples. Y=Yes, N=No, n/a=Not applicable. **(b)** Summary results for Patient 1 and Patient 2.

| Tool workflow* | CPU time (min) | Wall clock time (min) | Maximum resident set size (RAM in gigabytes) |
|---|---|---|---|
| NanoVar | 1186 | 194 | 31.7 |
| Picky | 16910 | 1194 | 25.1 |
| Sniffles | 13187 | 561 | 18.9 |
| NanoSV | 32076 | 5278 | 21.5 |
| Delly | 28281 | 3374 | 33.2 |
| novoBreak | 57157 | 3519 | 45.0 |

*Comprise of sequence mapping and SV calling, using 24 threads

**Supplementary Table 6**: Runtime and maximum memory usage consumed by the workflows of each tool using 24 threads for the SV characterization in Patient 1. For 3GS tools, 12 gigabases of sequencing data was used, while for 2GS tools, 160 gigabases of sequencing data was used. Data was collected using GNU Time.

| GenBank accession no. | Virus name |
|---|---|
| NC_012959.1 | Human adenovirus 54 |
| NC_001460.1 | Human adenovirus A |
| NC_011203.1 | Human adenovirus B1 |
| NC_011202.1 | Human adenovirus B2 |
| NC_001405.1 | Human adenovirus C |
| NC_010956.1 | Human adenovirus D |
| NC_003977.2 | Hepatitis B virus (strain ayw) |
| NC_001806.2 | Human herpesvirus 1 strain 17 |
| NC_001798.2 | Human herpesvirus 2 strain HG52 |
| NC_001348.1 | Human herpesvirus 3 |
| NC_007605.1 | Human herpesvirus 4 |
| NC_006273.2 | Human herpesvirus 5 strain Merlin |
| NC_001716.2 | Human herpesvirus 7 |
| NC_009333.1 | Human herpesvirus 8 |
| NC_017994.1 | Human papillomavirus type 136 |
| NC_017996.1 | Human papillomavirus type 140 |
| NC_021483.1 | Human papillomavirus type 154 isolate PV77 |
| NC_033781.1 | Human papillomavirus type 156 isolate GC01 |
| NC_001526.4 | Human papillomavirus type 16 |
| NC_023891.1 | Human papillomavirus type 178 |
| NC_022095.1 | Human papillomavirus type 179 |
| NC_001357.1 | Human papillomavirus 18 |
| NC_001356.1 | Human papillomavirus 1 |
| NC_027528.1 | Human papillomavirus type 201 isolate HPV201 |
| NC_001352.1 | Human papillomavirus 2 |
| NC_001591.1 | Human papillomavirus type 49 |
| NC_001531.1 | Human papillomavirus type 5 |
| NC_034616.1 | Human papillomavirus type 85 isolate 114B |
| NC_004500.1 | Human papillomavirus type 92 |
| NC_005134.2 | Human papillomavirus type 96 |
| NC_001596.1 | Human papillomavirus type 9 |
| NC_001401.2 | Adeno-associated virus 2 |
| NC_001729.1 | Adeno-associated virus 3 |
| NC_006152.1 | Adeno-associated virus 5 |
| NC_018102.1 | MW polyomavirus |
| NC_020106.1 | STL polyomavirus strain MA138 |
| NC_020890.1 | Human polyomavirus 12 strain hu1403 |
| NC_024118.1 | New Jersey polyomavirus-2013 isolate NJ-PyV-2013 |
| NC_001538.1 | BK polyomavirus |
| NC_001699.1 | JC polyomavirus |
| NC_009238.1 | KI polyomavirus Stockholm 60 |
| NC_009539.1 | WU Polyomavirus |
| NC_010277.2 | Merkel cell polyomavirus isolate R17b |
| NC_014406.1 | Human polyomavirus 6 |
| NC_014407.1 | Human polyomavirus 7 |
| NC_014361.1 | Trichodysplasia spinulosa-associated polyomavirus |
| NC_015150.1 | Human polyomavirus 9 |
| NC_001669.1 | Simian virus 40 |
| NC_022518.1 | Human endogenous retrovirus K113 |
| NC_001802.1 | Human immunodeficiency virus 1 |
| NC_001722.1 | Human immunodeficiency virus 2 |
| NC_001436.1 | Human T-lymphotropic virus 1 |
| NC_001488.1 | Human T-lymphotropic virus 2 |
| NC_001364.1 | Simian foamy virus |

**Supplementary Table 7**: GenBank accession number and name of viruses used for SV simulation (novel insertions).

| 3GS long-read | | | | | | | | | |
|---|---|---|---|---|---|---|---|---|---|
| Sample | Number of reads | Read length (bp) | | Total bases ($x10^6$) | Bases mapped ($x10^6$) | Percentage of bases mapped | Alignment error rate | Lengthwise genome coverage | Estimated sequencing depth |
| | | Median | Interquartile range | | | | | | |
| Simulation 1 | 2,000,000 | 6318 | 4372 | 12,800 | 12,690 | 99.14% | 1.70E-01 | 92.68% | 4.11x |
| Simulation 2 | 2,000,000 | 6316 | 4375 | 12,793 | 12,684 | 99.15% | 1.70E-01 | 92.67% | 4.11x |
| Simulation 3 | 2,000,000 | 6318 | 4375 | 12,797 | 12,687 | 99.14% | 1.70E-01 | 92.69% | 4.11x |
| Patient 1 | 1,793,667 | 6721 | 5226 | 12,407 | 11,662 | 93.99% | 1.65E-01 | 90.27% | 3.78x |
| Patient 2 | 1,939,432 | 6197 | 3798 | 12,304 | 11,583 | 94.14% | 1.86E-01 | 90.07% | 3.75x |

| 2GS short-read | | | | | | | | |
|---|---|---|---|---|---|---|---|---|
| Sample | Number of reads | Read length (bp) | Total bases ($x10^6$) | Bases mapped ($x10^6$) | Percentage of bases mapped | Alignment error rate | Lengthwise genome coverage | Estimated sequencing depth |
| Simulation 1 | 1,089,608,480 | 150 | 163,441 | 162,629 | 99.50% | 6.07E-03 | 94.49% | 52.66x |
| Simulation 2 | 1,089,649,172 | 150 | 163,447 | 162,633 | 99.50% | 6.08E-03 | 94.47% | 52.66x |
| Simulation 3 | 1,089,498,292 | 150 | 163,425 | 162,611 | 99.50% | 6.08E-03 | 94.47% | 52.65x |
| Patient 1 | 1,084,425,108 | 150 | 162,664 | 160,274 | 98.53% | 7.51E-03 | 94.52% | 51.90x |
| Patient 2 | 1,096,966,824 | 150 | 164,545 | 162,251 | 98.61% | 7.23E-03 | 94.51% | 52.54x |

**Supplementary Table 8**: Statistics of 3GS and 2GS reads and read mapping of actual reads and simulated reads. All reads were mapped using BWA aligner and statistics were calculated by SAMTools. The sequencing depth was estimated from the Lander/Waterman equation.

| SV id | Forward primer (5'-3') | Reverse primer (5'-3') |
|---|---|---|
| 1-1 | GCTCCAGTCAGATTGCACTAA | GCCTCCATAGACCTACAGTTTC |
| 1-2 | GAAAGGGAAGGTGATGCTGATA | AGCCTGTTTATGGTAATAGGGATAG |
| 1-3 | AGAACATTATTCCACCAGCCA | AGTGAGAAGAAGCAGAGAAATCA |
| 1-4 | GCTTGTAACAAACGGATGGG | CCTCTTAATTGCACTTTGACTCC |
| 1-5 | CAAAGTGTAAAGTGGCTTGGTG | GCTGGTAACCTTGTAGGGAATAG |
| 1-6 | GAGGCACAGACACATCCAAA | CTGCTTCAGCTTCTCTGTTCA |
| 1-7 | CAACCTCCTTGGTTAAATGTGTTC | CAGGATTAGTTGCTTGTTTCCTTAC |
| 1-8 | ATATGCCAGTCTGTGCTGTG | GCCCTGCCTATGCCTTA |
| 2-1 | AGAGACATCTCCAAACCTACAA | TCTGGGAAATCTGGCTCTAC |
| 2-2 | CATGGCTGTAAGTGGGTTTATC | CATATCGTTAGATTTCTTCACTGCT |
| 2-3 | CCCTAACTTCCAGGCTTCTA | AGTCACCTATGATCCTGCTTAT |
| 2-4 | CAAGGCAGAACCTCCAATG | ATCCTTGTGCCAGGTTATCT |
| 2-4 | *AGAGAGATCCAAGCTGTAGTC | *TTTGCATACACTGTGAAGTAGG |
| 2-4 | CAAGGCAGAACCTCCAATG | *GGCAGGTTTATATTTGTTTCAGTTC |
| 2-5 | GAGGCACAGGGACCATAAC | TGCCACAACAAATGGTTTACAA |
| 2-6 | AGTTGTAGCATCTACGTTAGACA | TGGGCATGAAGAGTGATAGG |
| 2-7 | GGGAATACTCTGAGATGTGTGA | CTGTATATCACTAGGATGTAGTAAAGTG |
| 2-8 | GCGTTGAGTTTCACAATGACTAA | GGGAGTTCTTTCACCTCTTTCT |

*Nested primers for reamplification of PCR product

**Supplementary Table 9**: Primers used for SV validation in Patient 1 and Patient 2 shown in Figure 3, and nested primers used for PCR product reamplification for Sanger sequencing (SV 2-4).
